# The expected behaviour of random fields in high dimensions: contradictions in the results of Bansal and Peterson (2018)

**DOI:** 10.1101/2021.01.21.427611

**Authors:** Samuel Davenport, Thomas E. Nichols

## Abstract

Bansal and Peterson (2018) found that in simple stationary Gaussian simulations Random Field Theory incorrectly estimates the number of clusters of a Gaussian field that lie above a threshold. Their results contradict the existing literature and appear to have arisen due to errors in their code. Using reproducible code we demonstrate that in their simulations Random Field Theory correctly predicts the expected number of clusters and therefore that many of their results are invalid.

---

The use of Random Field Theory (RFT) to control the false positive rate in neuroimaging is a well established testing framework. RFT consists of a set of theoretical results (originally due to Adler (1981)) that have been translated for use in neuroimaging (Worsley et al. (1992),Friston et al. (1994), Worsley et al. (1996)) and have been rigorously tested in the context of simulations (Hayasaka and Nichols (2003), Nichols and Hayasaka (2003). In a recent manuscript, Bansal and Peterson (2018) (henceforth BP) found that in simple stationary Gaussian simulations RFT incorrectly estimates the number of clusters of a Gaussian field that lie above a threshold. However, these results contradict the existing literature and our own previous findings, and exhibited highly counter-intuitive features. In particular, BP’s results showed the expected number of clusters falling then growing as a function of smoothness. Instead, as smoothness increases the expected cluster count should simply fall. BP’s authors helpfully provided some of their code and we identified some errors due to boundary wrap-around; the authors recomputed their results which then changed substantially but still did not align with previous results (i.e. there was still a great mismatch between theory and simulation). Unfortunately this implies that their conclusions regarding the performance of RFT are invalid.

Cluster size inference using Random Field Theory is based on the properties of null mean-zero homogeneous random fields thresholded at a given cluster defining threshold (CDT) to produce an excursion set (i.e. a collection of suprathreshold clusters). In this setting the expected Euler characteristic provides an approximation to the expected number of clusters. Then, under the assumption that clusters are elliptic paraboloids (true at high CDTs), results originally due to Nosko (1969), Wilson and Adler (1982) and Wilson (1988) provide a distribution for the extent of each cluster (see also Adler(1981) p158, Adler et al. (2010) chapter 6 and Cao (1999)). This was used to derive a form for the maximum cluster size above the CDT allowing control of the familywise error rate over clusters (Friston et al. (1994), Friston et al. (1996)). At low CDTs resting state data validation has shown that false positives are not correctly controlled using this approach (Eklund et al., 2016). They showed that this improves as the CDT is increased, in line with theory, and using high CDTs ensures good false positive rate control (Gong et al., 2018). However, RFT has long been known to perform well in simulations where the assumptions are satisfied.

The purpose of this note is to demonstrate with freely available and reproducible code the evaluations considered by BP; showing simulation and theory align. We find that RFT correctly estimates the expected number of clusters over a range of parameter settings. Analysis (random field simulation and thresholding) was performed using the RFTtoolbox, available at https://github.com/sjdavenport/RFTtoolbox, and SPM. Scripts to run this analysis are available at https://github.com/sjdavenport/BansalResponse. All code was run in MATLAB R2019b.

## 1 Methods

### 1.1 Random Field Generation

We generate Gaussian random fields (GRFs) using the RFTtoolbox. Given a number of dimensions *D* and a grid of size 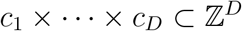 and a given FWHM *f*, this is done by generating i.i.d standard Gaussian random variables on a grid of size (*c*_1_ + 2*C*) × ··· × (*c_N_* + 2*C*) where *C* = ⌈1.7/*f*⌉ is a buffer to avoid boundary effects^1^. These are then smoothed with a Gaussian kernel with FWHM *f* and the central *c*_1_ × ··· × *c_D_* subset is taken as the output random field. Finally, the image is divided by the square root of the sum-of-squares of the kernel to ensure that the resulting field is unit-variance.

For our simulations we use the same settings as in BP. In particular for *D* = 1 we generate 1D GRFs consisting of 10000 voxels. For *D* = 2, we generate GRFs on a lattice of size 250 × 250 and for *D* = 3 we generate GRFs on a lattice of size 90 × 90 × 90. For each dimension, each CDT = 1.5, 2, 2.5, 3 and each FWHM = 5,10,15, 20, 25 voxels, we generate 100000 GRFs (an increase on the 50 used in each setting in BP). This allows us to compute the average (Monte Carlo) number of clusters above the threshold for each setting in order to compare to theoretical predictions.

### 1.2 Expected Number of Clusters

In the RFT framework, for large enough thresholds, the Euler characteristic of the excursion set is equal to the number of its connected components (Worsley et al., 1992). As such the expected Euler characteristic can be used to estimate the expected number of clusters above the threshold.

BP use the original formula (due to Adler (1981)) for the expected Euler characteristic. For a *D*-dimensional stationary random field *X* on a set 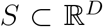 with standard deviation *σ*, a cluster defining threshold *u* and an excursion set *A_u_* above the threshold this gives

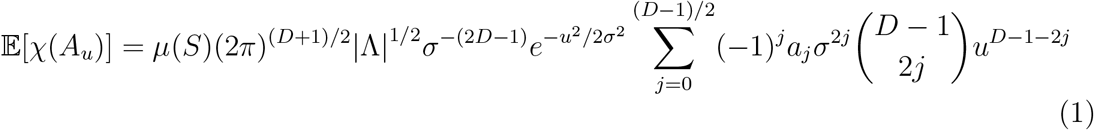

where Λ is the covariance matrix of the partial derivatives, *μ*(*S*) is the volume of *S* and *a_j_* = (2*j*)!/*j*!2^*j*^. Unfortunately BP appears to have miscalculated this (see Section 3). However this version is also out of date as it fails to account for the occurence of local maxima on the boundary. The updated formula due to (Taylor (2006)) is

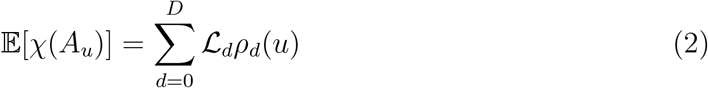

where for *d* = 1,…, *D*, *ρ_d_* are known functions (see Worsley et al. (1996) for their explicit forms) and 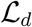 are known as the Lipshitz Killing curvatures (LKCs) which are determined by the covariance function of the random fields and the shape of the parameter set on which the fields are defined. When the random fields are stationary these can be calculated (Worsley et al., 1996) and this is the version of RFT that has been implemented in current software packages (such as SPM and FSL).

For stationary fields the LKCs are determined by Λ and S. In particular, for a *D*-dimensional field consisting of i.i.d Gaussian noise and smoothed with an isotropic Gaussian kernel with FWHM *s* (as in our and BP’s simulations), Λ is a diagonal matrix with diagonal entries Λ_*ii*_ = 4log(2)/*s*^2^ for *i* = 1,…,*D* (see Holmes (1994), Worsley et al. (1992)) and so the LKCs are known. Expansion (2) is in fact also valid under non-stationarity Taylor (2006). However in 3D the LKCs are rather difficult to compute and, while there has been promising recent work on this (Adler et al. (2017), Telschow et al. (2019)), it has not yet been implemented in neuroimaging software packages.

## 2 Results

Rerunning the simulations and theory correctly allows us to reproduce BP’s Table 1 in tables listed in Figure 4. We present these results for 2D and 3D in graphical form in Figures 1 and 2 where we plot the theoretical and average number of clusters. We show that there is large discrepancy between our results (which match previous work) and the results found by BP. The figures demonstrate that in these simulation settings RFT provides a close estimate of the expected Euler characteristic (at all applied smoothness levels) contradicting their findings.

**Figure 1:**
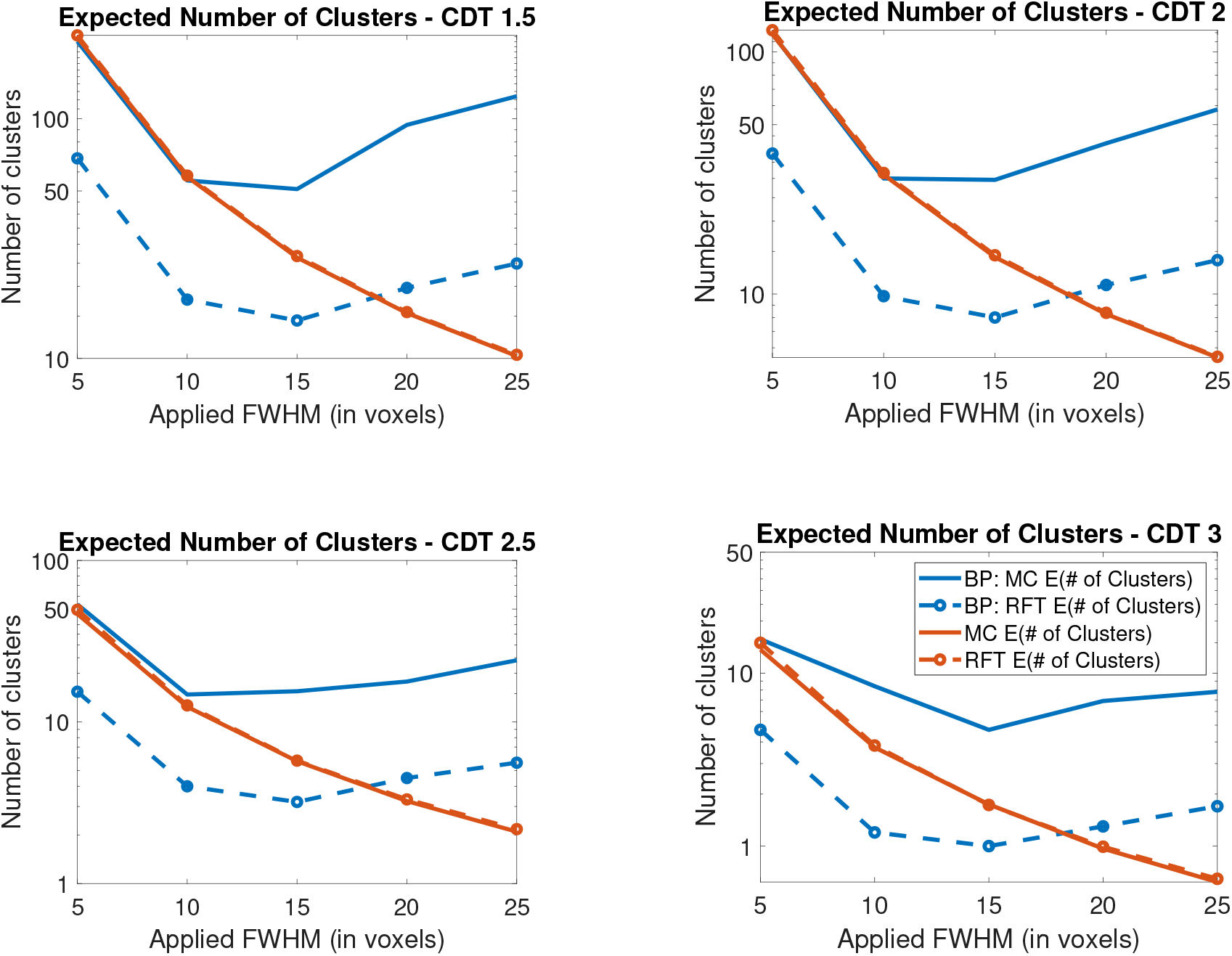
Comparison of Monte Carlo and theoretical expected number of clusters in 2D (in the legend # denotes the word number). In this setting the expected and average number of clusters (shown in red) closely match and are very different from those found in BP.

**Figure 2:**
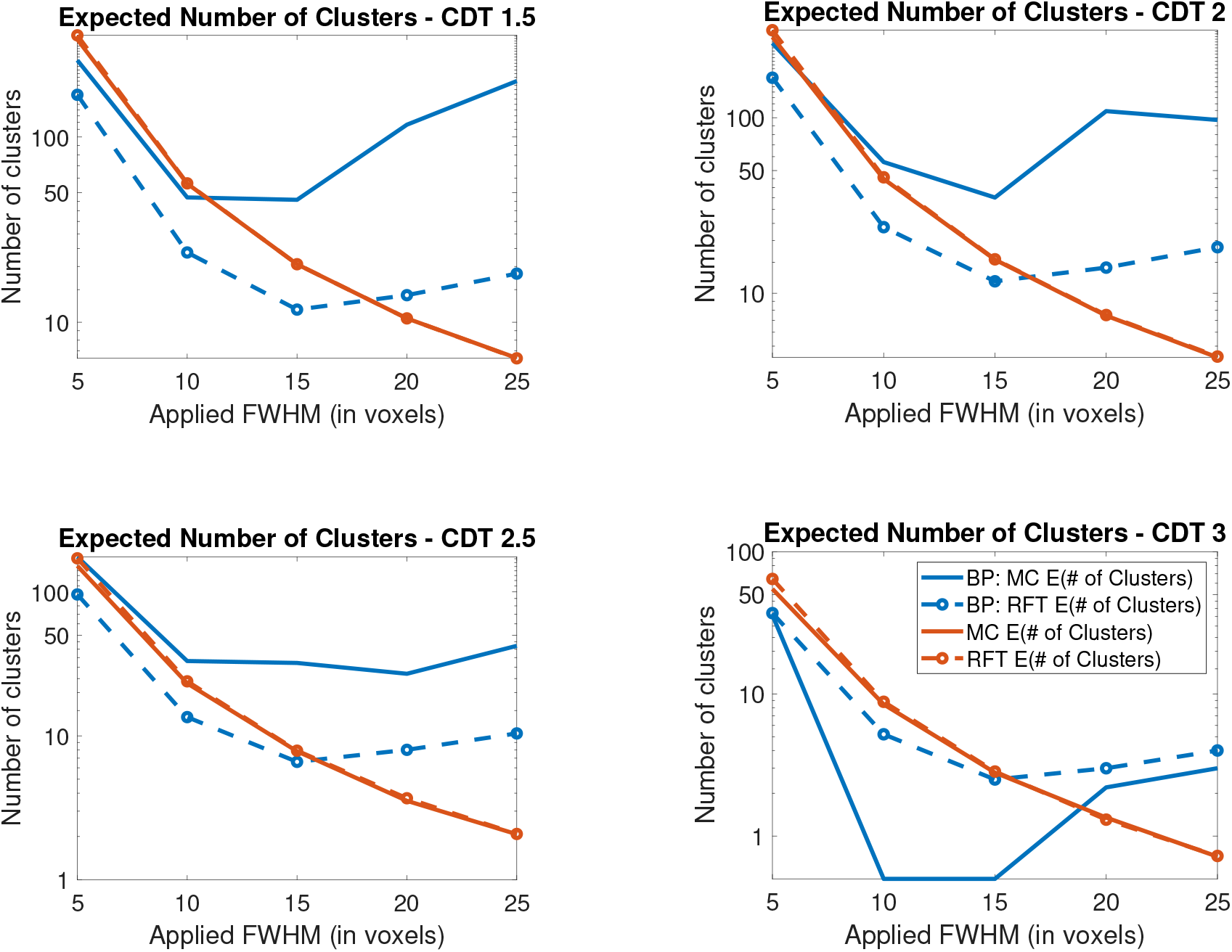
Comparison of Monte Carlo and theoretical expected number of clusters in 3D (in the legend # denotes the word number). In this setting the expected and average number of clusters (shown in red) closely match and are very different from those found in BP.

These figures additionally show that, as the smoothness increases, the number of clusters above the threshold decreases. The BP results instead show the number of clusters first decreasing and then increasing as the smoothness increases (this issue appears to affect both their calculations of the expected cluster size and of the average number of clusters above the threshold). For further details, and a comparison of the results, see the discussion.

The corresponding 1D results are shown graphically in Figure 3. Unlike the 2D and 3D results, in this setting, at low CDTs BP’s results closely match ours. However, as the CDT is increased, their calculation of the average number of clusters no longer matches. This could partially be due to noise as they only used 50 simulations in each setting.

**Figure 3:**
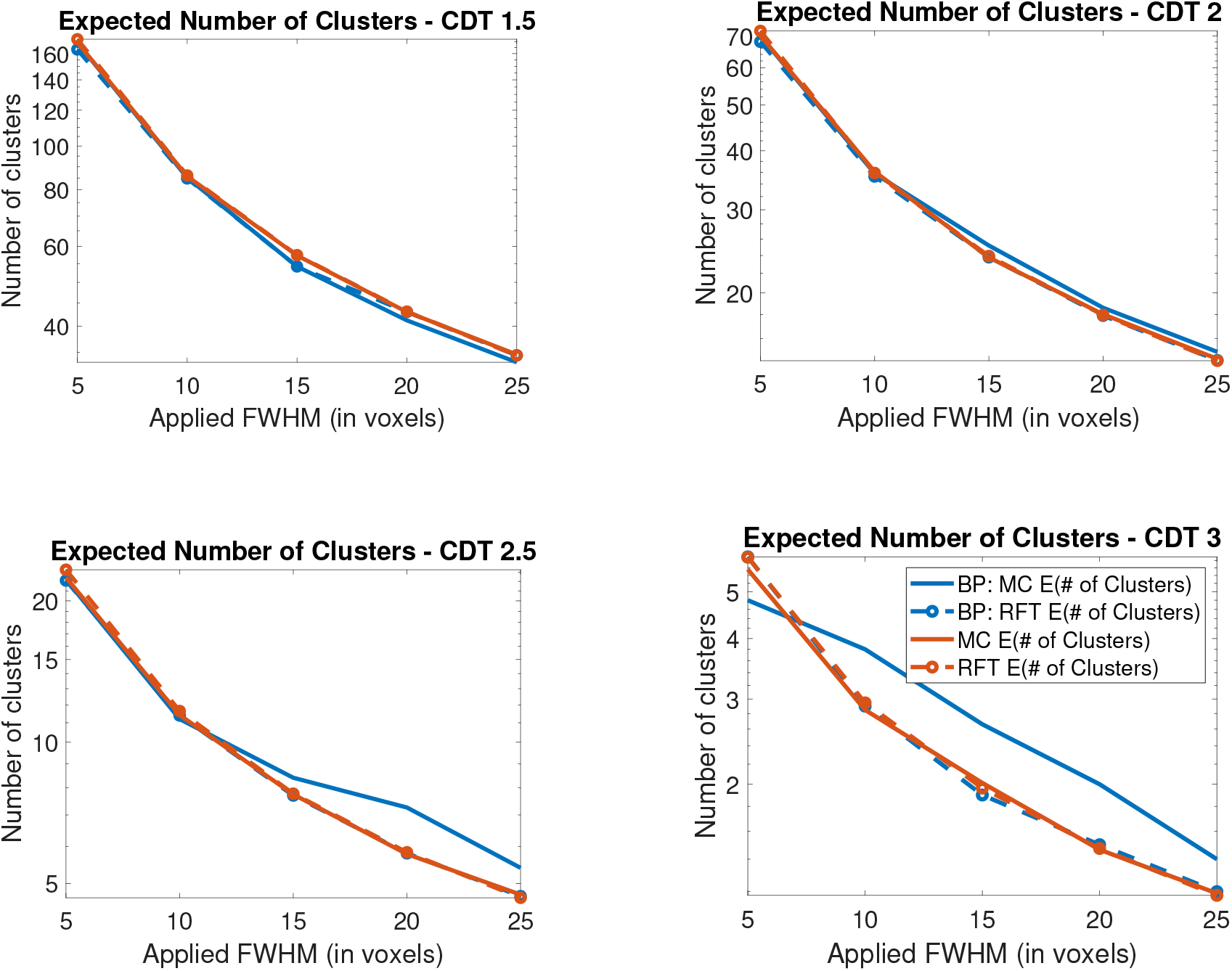
Comparison of Monte Carlo and theoretical expected number of clusters in 1D (in the legend # denotes the word number). At low CDTs our results match those of BP, however at high CDTs the average number of clusters that they find is slightly inflated but the theoretical number of clusters they find matches our calculations.

## 3 Discussion

Our results show that in these simulations the expected Euler characteristic closely matches the average number of clusters as predicted by theory. Unfortunately it appears that BP have miscalculated the Euler characteristics and have simulated their random fields incorrectly. Their work predicts large differences between theory and simulations but this is unfounded. We have regenerated corrected versions of their Table (Figure 4) and have presented these graphically. Our results are very different from those of BP and show that the expected Euler characteristic clearly matches the average number of clusters, providing a validation of the theory in line with previous work (e.g. Hayasaka and Nichols (2003), Worsley et al. (1996)).

**Figure 4:**
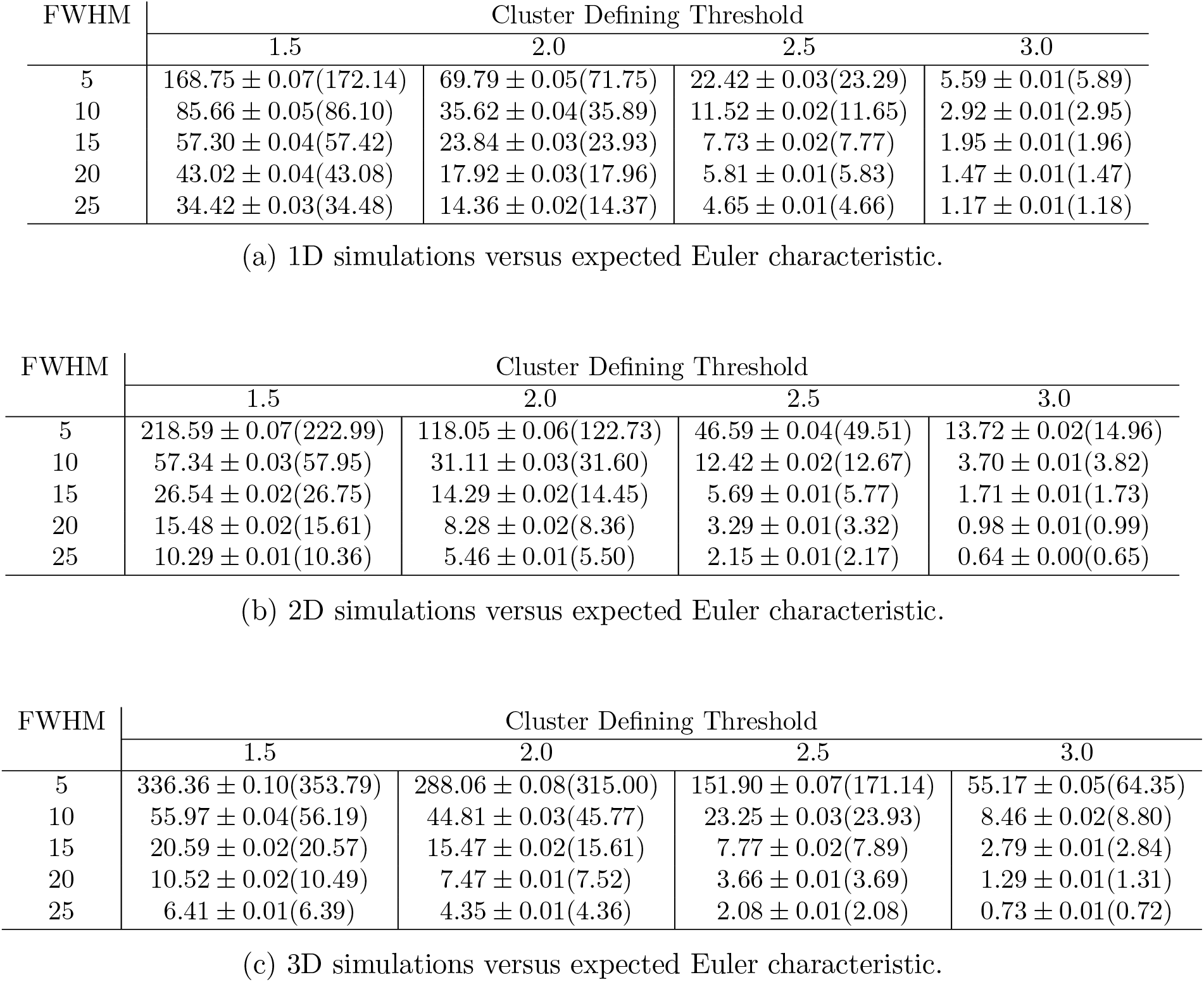
Comparing the average number of clusters above a threshold with the expected Euler characteristic. For each table each entry is of the form *A* ± *s*(EEC) where *A* is the average number of clusters above the threshold, ±*s* provides a 95% confidence interval for this average and EEC is the expected Euler characteristic provided by theory. This shows that the EEC provides a good approximation of the expected number of clusters above the threshold.

To immediately see that their results are incorrect, note the non-monotonicity of the blue curves shown in Figures 1 and 2. This non-monotonicity is present in both their theoretical and their Monte-Carlo calculations. However this indicates a significant error in their calculations because as the applied smoothing increases the number of clusters above the threshold should decrease. Indeed one only needs to study Equation (1), to see that as the FWHM increases the expected Euler characteristic should decrease.

It is difficult to identify the precise source of the error in the results of BP. One issue is that they use Equation 1 (which does not account for boundary effects) to calculate the expected Euler characteristic instead of the updated formula (2) which is implemented (under stationarity) in neuroimaging software. Another source of error can be seen by examining Figure 1 from their paper. Close study of this Figure suggests that the data has been simulated on a 2D torus rather than on a grid as the clusters on the edge of the image can be matched (in correspondence with the Authors we confirmed that this is how they simulated their random fields). This could contribute to the non-monotonicity because clusters at the boundary would be counted twice and boundary clusters are more likely to occur at higher smoothness levels. However, this setting is not representative of the datasets used in fMRI and in particular equations (1) and (2) for the expected Euler characteristic of the excursion set are only valid for a closed and bounded subset of 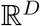 and not for the torus. The theory for such a space would be completely different and so these formulae cannot be applied. This is thus a partial explanation for the disagreement (as their 3D simulations suffer from a similar problem). However even accounting for that we were not able to replicate the non-monotonicity so it seems as though there may be deeper problems with their code. Unfortunately, as a result, many of the conclusions of their paper are likely invalid. We suspect that there are a number of further underlying issues with their code since, for instance, they also appear to have miscalculated the expected Euler characteristics (as these also suffer from non-monotonicity in their results).

BP appear to have incorrectly generated their random fields and to have miscalculated the expected Euler characteristic, meaning that their conclusions should not be relied upon. We have shown that their results disagree with our simulations which supported the existing well-established and validated theory.

## 4 Appendix

## 5 Acknowledgements

We would like to thank the 2 anonymous reviewers for their comments which helped to improve the quality of the manuscript.

TEN is supported by the Wellcome Trust, 100309/Z/12/Z and SD is funded by the EPSRC.

## Footnotes

1 For 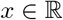 denotes the smallest integer than is greater than or equal to *x*.

## Notes

https://github.com/sjdavenport/RFTtoolbox

### Competing Interest Statement

The authors have declared no competing interest.

